# The HERV-K(HML-2) Transcriptome in Non-Diseased Tissue

**DOI:** 10.1101/2022.06.24.497521

**Authors:** Aidan Burn, Farrah Roy, Michael Freeman, John M Coffin

## Abstract

Human endogenous retrovirus (HERV) transcripts are known to be highly expressed in cancers, yet their activity in non-diseased tissue is largely unknown. Using the GTEx RNA-seq dataset from normal tissue sampled at autopsy, we characterized individual expression of the recent HERV-K(HML-2) provirus group across 13,000 different samples of 54 different tissues from 948 individuals. HML-2 transcripts could be identified in every tissue sampled, and were elevated in the cerebellum, pituitary, testis, and thyroid. 37 different individual proviruses were expressed in one or more tissues, representing all three LTR5 subgroups. 9 proviruses were identified as having LTR-driven transcription, 7 of which belonged to the most recent LTR5HS subgroup. Proviruses of different subgroups displayed a bias in tissue expression which may be associated with differences in transcription factor binding sites in their LTRs. Provirus expression was greater in evolutionarily older proviruses with an earliest shared ancestor of gorilla or older. HML-2 expression was significantly affected by biological sex in one tissue, while age and timing of death (Hardy score) had little effect. Proviruses containing intact *gag, pro* and *env* ORFs were expressed in the dataset, with almost every tissue measured potentially expressing at least one intact ORF (*gag*).

## Introduction

Retroviruses have been infecting mammals and other vertebrates for at least 100 million years, invading somatic and germ cells of their host species (Bannert & Kurth, 2006). Proviral remnants of ancient retroviral infection of germ cells now make up about 8% of the human genome in the form of human endogenous retroviruses (HERVs)(Lander et al., 2001; Li et al., 2001). These ancient retroviruses infected the germline of ancestral primates, and the resulting proviruses are inherited in a Mendelian fashion and subject to fixation or loss over time. Many of them have become significantly degraded, leaving behind remnants in the form of solo **L**ong **T**erminal **R**epeats (LTRs), and fragments of viral **O**pen **R**eading **F**rames (ORFs). Regardless, their presence in the primate genome allows one to piece together the history of retroviral infection in primates, a process analogous to use of partial fossil remains of multiple individuals of a species to understand its evolutionary history.

There are at least 30 HERV subclades. Many are distantly related to exogenous viruses extant today in other animals; all represent apparently extinct retroviral lineages. Of all the HERV groups the only subclade containing human-specific - and therefore the most recently integrated - proviruses can be found within the **H**uman **M**ouse mammary Tumor Virus **L**ike (HML)-2 subgroup within the HERV-K group belonging to the *Betaretrovirus* genus. The first record of its integration dates to ∼40 million years ago, continuing until as recently as 200,000 years ago (Hanke et al., 2016; Nelson et al., 2003; Turner et al., 2001). HERV-K (HML-2), hereafter referred to as “HML-2”, has undergone multiple bursts of integration into the human ancestral germline leaving a number of proviruses that contain intact ORFS and polymorphic insertions across the human population (Figure 1A) (Turner et al., 2001). The HERV-K group is named for its inferred lysine tRNA primer utilized for initiating reverse transcription (Larsson et al., 1989). The LTR regions of each HML-2 provirus can be used to group the provirus phylogenetically into 3 main subgroups; LTR5A, LTR5B and LTR5HS. The LTR5HS subgroup generally consists of more recent integrations, while the LTR5A and LTR5B subgroups are thought have been active before human speciation (Buzdin et al., 2003; Subramanian et al., 2011).

**Figure 1.**
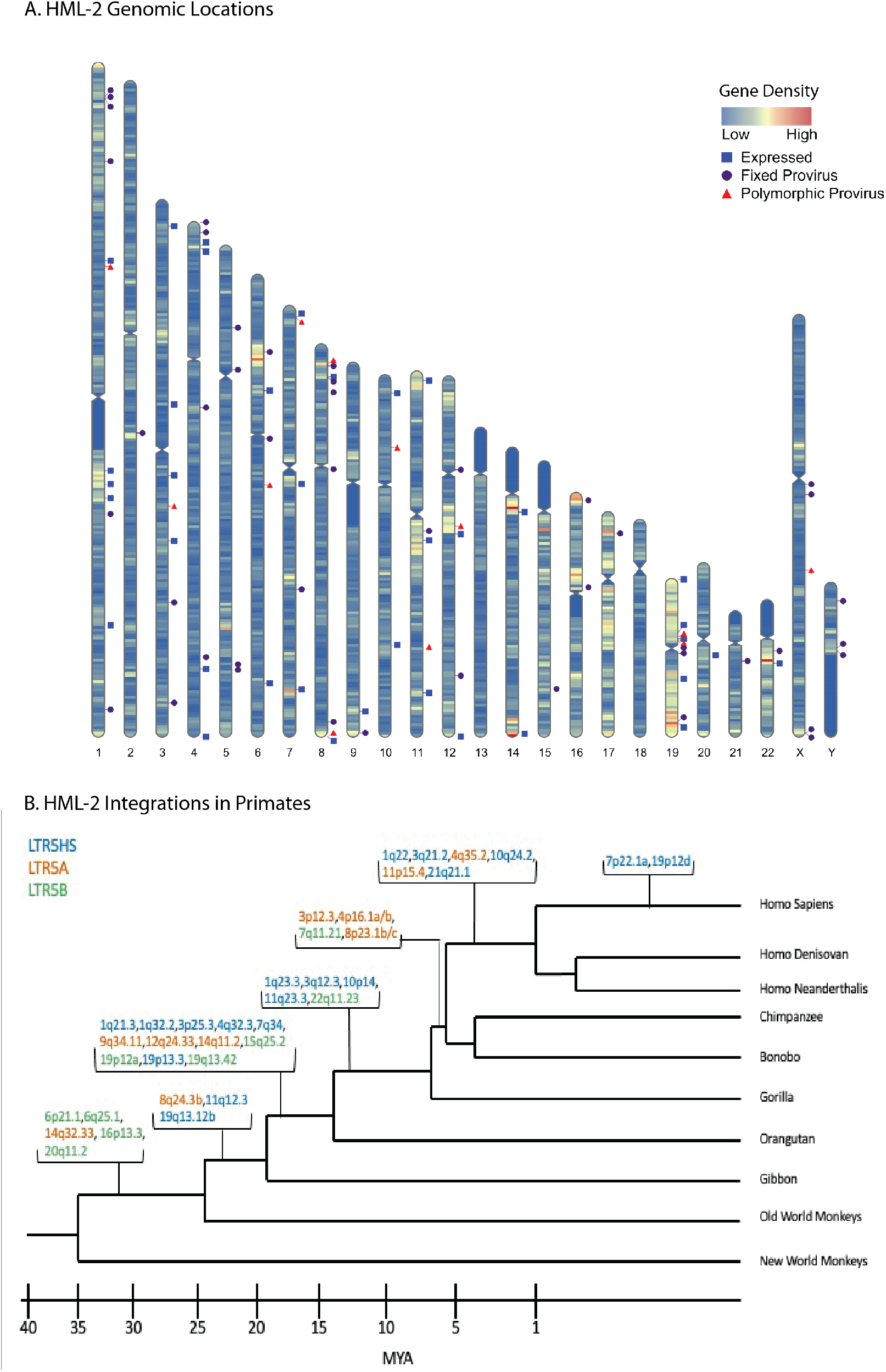
HERV-K HML-2 proviruses vary in chromosomal location and age of integration. A. Human chromosome map showing gene density and location of 2-LTR proviruses in the genome. Purple circles represent proviruses that are fixed in the human population; red triangles represent known polymorphic proviruses, blue squares represent GTEx expressed (TPM ≥1) proviruses. Note that the expression status of polymorphic proviruses could not be determined. Gene density is represented in a heatmap of low = blue to high = red. B. Primate phylogeny illustrating the earliest common ancestor of HML-2 proviruses expressed in the GTEx dataset at a TPM of ≥1. As in this and subsequent figures, provirus color (blue, red, green) corresponds to LTR subtype (LTR5HS, LTR5A, and LTR 5B, respectively).

HML-2 proviruses have accumulated insertions, deletions, and internal recombination events over time, leaving defective remnants to be enriched by selection against their potentially pathogenic ancestors. Consequently, all annotated HML-2 proviruses in the human genome are defective for replication. There are currently 94 HML-2 proviruses that retain some internal sequence - sometimes referred to as “full length” - within the hg38 human reference genome along with at least 944 solo LTRs (Subramanian et al., 2011; Wildschutte et al., 2016). We will refer to the former group as “2-LTR” proviruses. Most human HML-2 proviruses are also found at the orthologous site in chimpanzees, implying that they are >5 million years old but around 26 are human specific (implying a younger age) and at least 14 of them remain un-fixed in the human population (Turner et al., 2001; Wildschutte et al., 2016). These relatively young proviruses have been subject to comparatively little evolutionary pressure, unlike the older proviruses, and therefore some retain at least one intact ORF (Dewannieux et al., 2005).

As illustrated in Figure 1A, HML-2 proviruses can be found throughout the human genome and are enriched in regions of high gene density. This insertion pattern reflects the preference of ancestral exogenous virus integrase for integration into areas of high gene expression (Brady et al., 2009), but insertions within genes are often subject to negative selection due to deleterious effects on their host (van de Lagemaat et al., 2006).The resulting variety of integration sites leads to dramatic differences in the surrounding genomic neighborhood from one provirus to the next. 60% of HML-2 insertions are within 30 kB of genes and 20% of all HML-2 proviruses are within introns, 80% of which are antisense to the surrounding gene (Subramanian et al., 2011).

While no HERV provirus has been shown to retain infectivity, an integrated provirus can affect the host in multiple ways (Subramanian et al., 2011). For example, genes encoding Env proteins of some ERV groups have been co-opted in mammals multiple times. The most famous example are ERV encoded Env proteins, now referred to as *syncytins* (Dupressoir et al., 2012). The cell-to-cell fusogenic ability of syncytins plays an important role in the formation of the syncytiotrophoblast during pregnancy in placental mammals (Lavialle et al., 2013). In a remarkable example of convergent evolution, different ERVs play the same role in most or all other placental mammalian orders (Dupressoir et al., 2012). Perhaps the most important evolutionary effect is that, in some cases, expression of modified ERV Env or Gag proteins can interfere with the replication of related exogenous viruses by blocking access to a receptor or interfering with capsid assembly or intracellular trafficking. Many well studied examples of ERV-mediated interference have been found in chickens, mice, cats, sheep, and other species (Best et al., 1996; Ito et al., 2016; Ito et al., 2013; Murcia et al., 2007; Yan et al., 2009). Notably, a co-opted HERV-T *env* gene is thought to have led to the extinction of the cognate exogenous virus in our primate ancestors (Blanco-Melo et al., 2017)

Proviruses with intact protein coding genes may be capable of expressing these genes upon activation in pathological states such as cancer. Some cancer cells and cell lines express the HML-2 structural protein Gag and can form mature viral particles (Bhardwaj et al., 2015; Löwer et al., 1984; Montesion et al., 2018a), and anti-Gag antibodies have been reported in individuals suffering from multiple malignancies such as breast cancer, melanoma, and teratocarcinoma (Hohn et al., 2013; Ruprecht et al., 2008). Another viral structural protein of disease-relevant interest is Env, a functional gene for which has been retained by at least one HML-2 provirus in the human genome, known as 7p22.1a or K108L. This protein retains its original fusogenic capacity, possibly allowing for formation of syncytia upon expression and has been found to be expressed in cancerous tissue and cell lines (Bhardwaj & Coffin, 2014; Dewannieux et al., 2005; Huang et al., 2013). The accessory protein Rec, which is required for unspliced viral RNA transport, has been shown to interact with a tumor suppressor in germ cell tumors and its expression is linked to a number of malignancies, although no causal relationship has yet been defined (Ruprecht et al., 2008). A subset comprising about half of the two-LTR HML-2 proviruses, termed “Type 1,” shares an identical 292 nucleotide deletion affecting the pol-env border. This deletion removes a splice donor for Rec, creating the transcript for an ORF called NP9. When translated, the NP9 protein has also been shown to interact with and interfere with a protein in the Numb/Notch pathway in germ cell tumors (Ruprecht et al., 2008). The type 1 deletion is found in proviruses shared by humans, chimpanzees and gorillas, implying that it arose more than 10 million years ago and is found only in LTR5Hs proviruses.

A provirus can also have significant effects on the host without viral gene expression. Proviruses introduced into the genome carry along with them promoters and poly(A) addition sites, as well as multiple splice acceptors. These sites can create alternative or intergenic splicing among host genes, altering transcription and affecting the integrity of the final protein product (Ruprecht et al., 2008). Binding sites for different transcription factors within proviral LTR promoters can greatly affect surrounding gene expression. These LTRs can act as promoters or enhancers both while flanking or within the integrated provirus or as solo LTRs near transcription start sites. The variety of binding sites in an LTR can affect neighboring gene expression in many ways, including direct changes to tissue specificity or by enhancing the activity of an existing promoter, depending on the location of integration and the surrounding genes (Feuchter & Mager, 1990; Jern & Coffin, 2008; Xiang et al., 2022).

Despite the potential importance of HML-2-modified expression of viral or host genes, to date, HML-2 expression in a non-diseased human body has not been well characterized. Data on HML-2 expression exists in various disease contexts, along with evidence of expression in some non-diseased tissues (Hohn et al., 2013; Schmitt et al., 2015), but examination of all tissues in a host has yet to be performed. Furthermore, most studies analyzing provirus expression in disease and non-disease context looked at total HML-2 expression rather than provirus-specific expression. Reports of total HML-2 expression fail to capture the diversity of HML-2 proviruses in the human genome and the variety of mechanisms controlling their individual expression. To create a detailed characterization, we turned to the **G**enotype **T**issue and **Ex**pression (GTEx) Project (Consortium, 2020). The GTEx project is a database of tissue specific gene expression collected from 54 non-diseased tissues across 948 donors. The wealth of RNA-Seq data contained in this database allows us to characterize HML-2 expression across the entire human body. By analyzing these data, we were able to discover evidence of HML-2 provirus expression in every tissue analyzed, with numerous proviruses showing significant expression across the body. The cerebellum, pituitary, testis, and thyroid showed the highest level of HML-2 expression in non-diseased tissue. A number of proviruses with intact ORFS were found to be expressed, although the functional consequences of their expression remain unclear. Common covariates such as biological sex showed significant differences between individual provirus expression on the basis of biological sex but not in other covariate groups. Furthermore, some proviruses appear to be self-driven by their 5’ LTRs, while many appear to be transcribed through other means, such as read-through transcription, either from a nearby gene or from an unannotated feature within the genome. Our study also revealed interesting patterns of expression, in that the oldest proviruses are the most expressed and most frequent provirus expression was seen in neuronal, endocrine, and reproductive tissue.

## Results

### Specific tissues have higher abundance of HERV-K(HML-2) Transcripts

Aberrant HML-2 expression in human cancer and a number of other disease states was recognized many years ago (Bhardwaj et al., 2015; Hohn et al., 2013; Löwer et al., 1984; Ruprecht et al., 2008), and its potential use as a biomarker for detection of illness or as therapeutic targets has long been discussed (Gimenez-Orenga & Oltra, 2021; Laumont et al., 2018; Tavakolian et al., 2021). In addition to their clinical use, there is a great deal of evolutionary biology one can learn from studying the genetic structure and expression of these ancient viral proviruses. For example, the tissue specificity of expression of individual proviruses can shed light on the mechanisms of replication, transmission, and pathogenesis of the ancestral virus. Taking advantage of the recently released human Genotype-Tissue Expression (GTEx) dataset (Consortium, 2020), we sought to profile HML-2 provirus expression in healthy tissues (Figure 2). This dataset is the most comprehensive listing of RNA-seq data for human tissues, consisting of 13,851 samples across 54 different tissues in the body acquired from 948 different postmortem donors. Reads were quality trimmed (Phred score > 30, min length of 75) before alignment to hg38 using HISAT2 (Kim et al., 2019). Aligned reads were then counted using the Bayesian analysis routine of the Telescope program (Bendall et al., 2019), which aligns multi-mapping reads to the most likely location based on a statistical model. Ambiguous reads were mapped through an iterative process that compares initial weights of read alignments to expected transcript levels. This procedure allows for more confident alignments of multi mapping reads for elements, like endogenous retroviruses, which are composed of repeated sequences that differ only slightly from one another.

**Figure 2.**
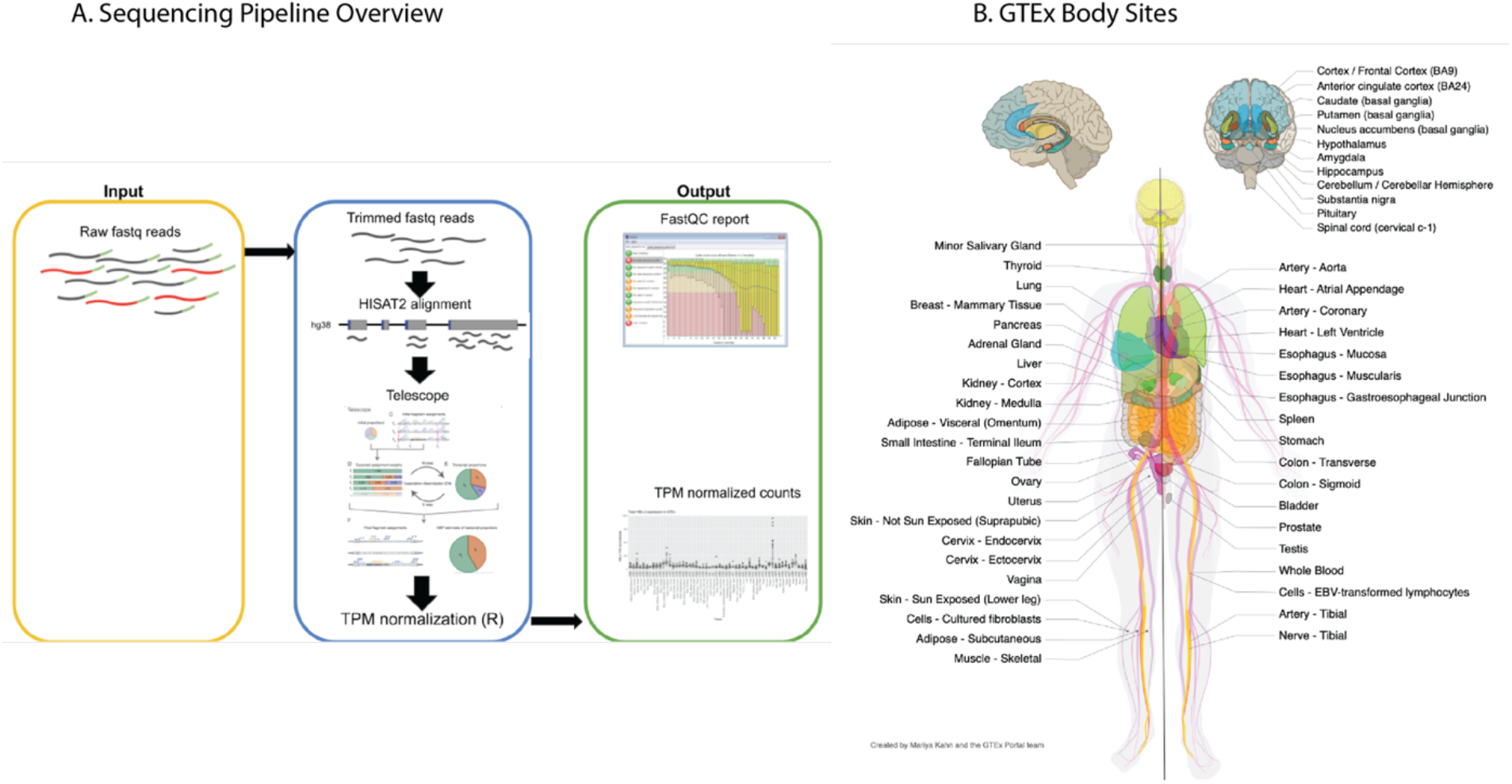
Analysis of the GTEx dataset A. Schematic overview of the sequencing pipeline used in this study. See Materials and Methods for further information. B. Diagram of all body sites samples in the GTEx project. Not listed is one cell culture, a chronic myelogenous leukemia line derived from a GTEx donor.

Our approach detected HML-2 expression throughout the body, with identifiable transcript levels (threshold ≥ 1 TPM) found in each body site analyzed (Figure 3A). The influence of the site sampled was immediately clear, with total HML-2 expression levels varying considerably across the body. Four tissues - cerebellum, pituitary, testis, and thyroid - had relatively high levels of HML-2 expression in comparison to other measured tissues within the GTEx cohort: these five tissues expressed combined total HML-2 transcripts at an average of at least 70 TPM across all samples. In contrast, samples from pancreas and whole blood showed lower detectable total HML-2 expression of 5 and 8 TPM/sample, respectively. In addition to the cerebellum, the rest of the brain and other nervous tissues appeared to be sites of relatively high levels of provirus expression, while tissues related to the circulatory or digestive systems showed lower levels of expression (apart from the spleen). Reproductive tissue such as the testis and ovaries also showed elevated expression compared to other tissues. The variation in HML-2 expression observed among these tissues demonstrates that the intensity of total HML-2 expression is tissue specific with some tissues and organ systems promoting expression more than others. Variation was also observed on an inter-donor level. While these figures represent data from all 948 donors, comparison of two individual donors shows another level of variability (Supplemental Figure 1). Provirus expression levels, specific provirus allele, and the number of proviruses expressed can change from donor to donor. This variability may result from a number of experimental factors, such as variable collection of tissue samples or could represent biological differences between the donors themselves, such as different health conditions, age, biological sex, or tissue condition at the time of collection. We examined these variables in subsequent analyses.

**Figure 3.**
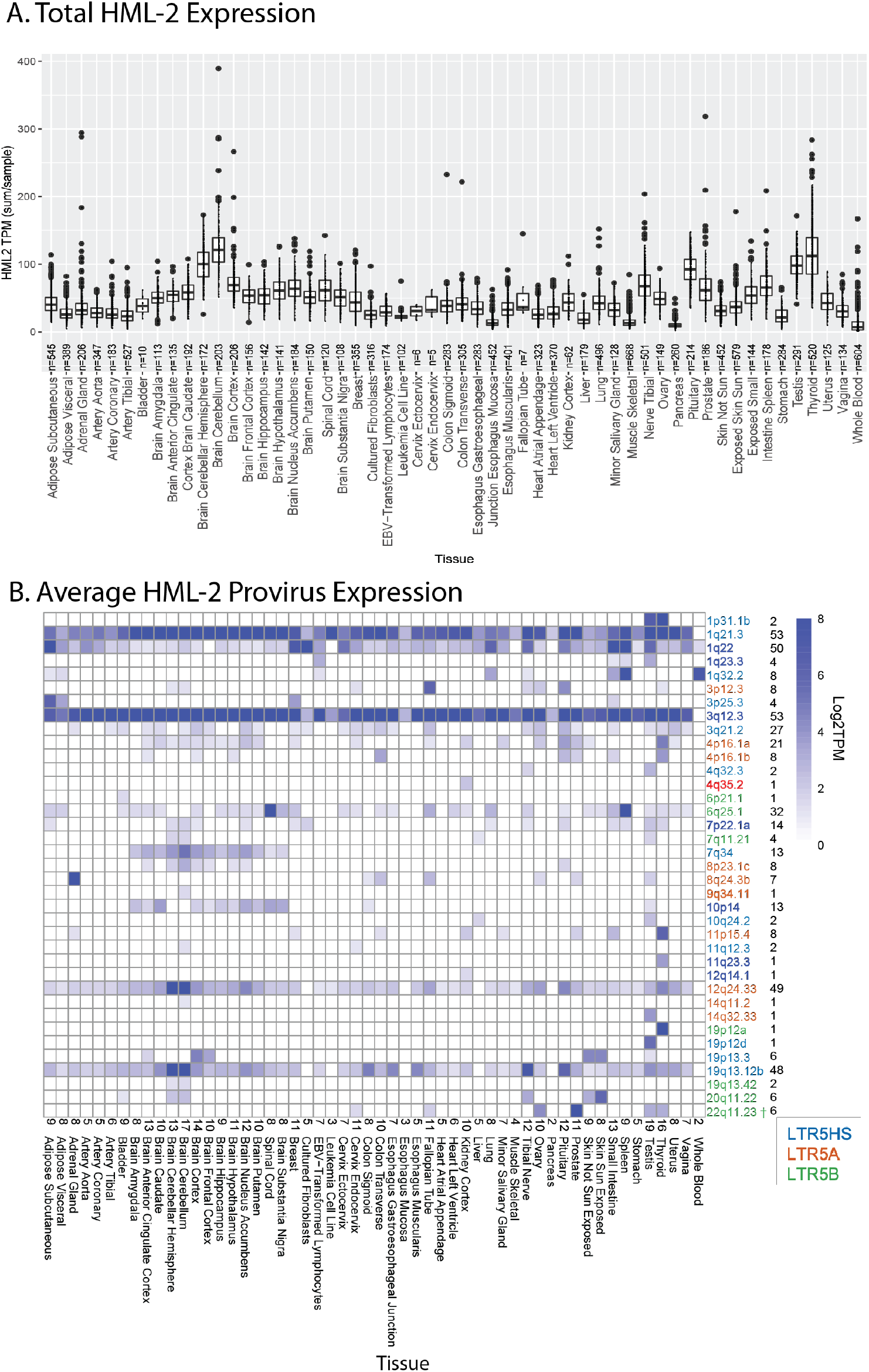
HML-2 Expression in GTEx. A. Combined expression of all HML-2 proviruses per individual sample per body site. The box and whisker plots show the mean as a line in the middle of a box bounded by the first and third quartile values, with the whiskers extending to 1.5 times the IQR (interquartile range) and with outlier values shown as individual dots. All proviruses with an average expression less than 1 TPM are excluded. B. Overall expression in log_2_ TPM for HML-2 proviruses expressed at ≥ 1 TPM is displayed for each body site. The specific proviruses, color coded by LTR type as in Figure 2, are shown on the right, followed by the number of tissues where each provirus was expressed. The number of proviruses in each tissue is displayed above the tissues on the horizontal axis. LTR driven proviruses are in **boldface**. † denotes the provirus 22q11.23 which is driven by a separate upstream 5HS LTR and was therefore not bolded in this figure.

While the total transcriptional level of HML-2s was of interest, it is only a small part of the picture. Although the HML-2 group comprises a set of closely related proviruses, each provirus is unique and is defined by individual mutational variation along with a unique genomic neighborhood that may influence or be influenced by its transcription. Therefore, exploring HML-2 expression on the individual provirus level is critical for understanding its evolution, functional consequences, and potential therapeutic impact. To properly investigate HML-2 expression in non-diseased tissue, the total HML-2 expression was broken down to the expression of each HML-2 provirus within each tissue with the aid of Telescope, allowing us to use partial expression data to accurately reconstruct the expression level, structure, and genomic location of each provirus (Bendall et al., 2019).

We detected 37 HML-2 proviruses expressed at levels above 1 TPM in at least one tissue. Individual provirus transcription could be classified into one of three different patterns of expression (Figure 3B): 1. Those that were expressed in almost all tissues sampled (apart from whole blood); 2. Those for which transcripts were only seen in tissues of a specific type; 3. Those that were transcribed in various tissues from multiple types. There were five proviruses that demonstrated the first pattern: 1q21.3, 1q22, 3q12.3, 12q24.33 and 19q13.12b. 1q21.3 and 3q12.3 were the highest expressed with levels regularly exceeding 8 TPM. The second pattern included proviruses, such as 10p14, 4p16.1a, and 7q34, which were expressed almost exclusively in the brain samples. The remainder of the expressed proviruses seen in the GTEx dataset, following the third pattern, were only expressed in a small number of tissues, and did not appear to be broadly associated with specific tissue types or organ systems. We observed high levels of HML-2 activity in prostate tissue, largely consisting of transcripts from 22q11.23, with lesser contribution from 1q21.3 and 3q12.3. Overall, it is clear that each tissue in the human body has a unique provirus expression pattern, and apart from the two proviruses expressed in 53/54 tissues, each provirus exhibited a unique pattern of expression.

### Mechanisms of HML-2 expression

As described in the introduction, proviruses are canonically expressed using enhancer and promoter signals in the 5’ LTR to initiate transcription at the U3-R border. However, as we reported previously (Bhardwaj et al., 2015; Montesion et al., 2018a), other mechanisms, including read-through from nearby or surrounding host genes, can also be used, particularly in cases where the 5’ LTR has been damaged by mutation. Examination of the alignments of RNA-seq reads from the GTEx data using Integrated Genomics Viewer (Thorvaldsdottir et al., 2013) can aid in elucidating such mechanisms. Using this method, the transcriptional mechanism of each provirus was classified (Supplemental Table 1). It is expected that for an LTR-driven provirus the aligned reads would map to the presumptive transcription start and poly(A) sites at the U3-R and R-U5 borders in the LTRs Figure 4A, for example, shows the data for the provirus at 3q12.3, one of the highest and most broadly expressed proviruses in the dataset. These aligned read clusters appear to start at the immediate 5’ end of the provirus, rather than the expected U3-R border, but this is likely an artifact resulting from double mapping of reads from the 3’ LTR. 9 HML-2 proviruses expressed in GTEx displayed an LTR driven mechanism. Reads were clearly observed to align at the TSS without aligned reads preceding the 5’ LTR. Each of these proviruses contain an intact 5’ LTR capable of driving transcription. All but two of these proviruses, those at 22q11.23 and 4q35.2, are LTR5HS proviruses. The provirus at 22q11.23 is an LTR5B provirus, yet transcription is driven by an LTR5HS promoter 551 bp upstream. This result was previously observed in the Tera-1 Teratocarcinoma cell line (Bhardwaj et al., 2015) and was seen again in our GTEx data (Supplemental Figure 2). 4q35.2 is an LTR5A subtype provirus, and the only one apparently expressed from its own LTR.

**Figure 4.**
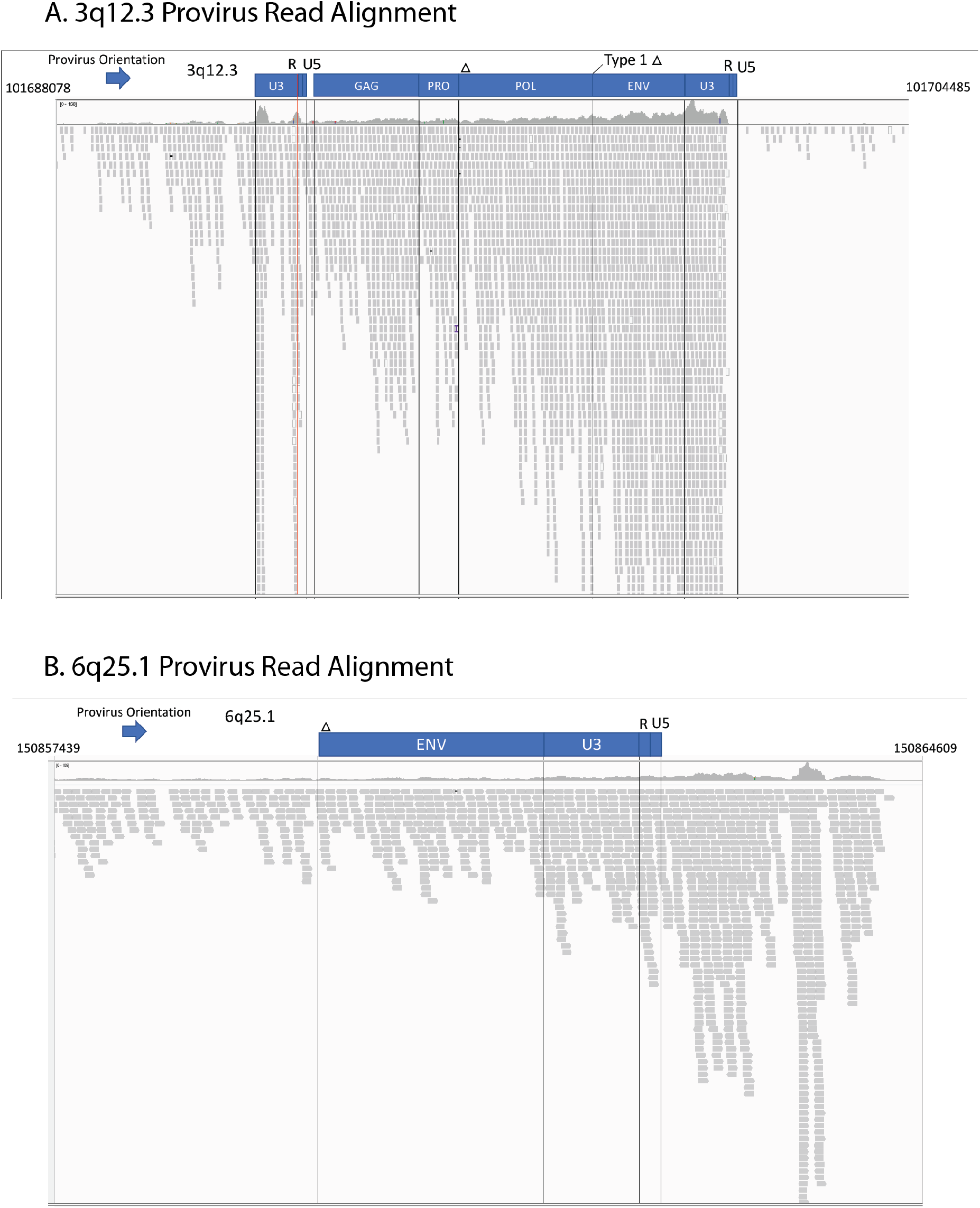
Expression of HML-2 proviruses can be self-driven or a result of the surrounding genomic neighborhood. These Integrated Gene Viewer snapshots show the alignment of GTEX RNA-seq reads to two proviruses. A. 3q12.3 This representative alignment is from the cerebellum of one GTEx donor. The red line marks the location of the transcription start site. B. Alignment of 6p25.1 from the spleen of another GTEX donor. Vertical lines demarcate divisions between the viral ORFs. Blue boxes denote provirus and gene structure. Genomic coordinates are displayed on the top left and right corners. The arrow denotes the orientation of each provirus.

Unlike these 9 LTR driven proviruses there were 24 expressed proviruses that either did not demonstrate clear LTR driven transcription starting at the 5’ LTR or did not contain a 5’ LTR at all. The provirus at 6q25.1 provides an example of such readthrough transcription (Figure 4B). This provirus completely lacks a 5’ LTR and adjacent *gag-pro-pol sequence*, yet the entirety of the remaining ∼2,800 sequence is expressed. Aligned reads can be seen stretching from upstream of the provirus into the provirus sequence with no changes or characteristic gaps suggesting a new TSS. This even alignment of reads combined with the lack of a required LTR sequence suggests that expression of this provirus sequence is driven by an outside element. Interrogating the source of RNA-seq reads is complicated by technical limitations of the GTEx dataset. Longer read lengths and stranded sequencing would provide a clearer picture of the direction of expression, as well as the clear start and end and splicing pattern of provirus transcripts. Yet it is clear that there are multiple mechanisms driving provirus expression in non-diseased human tissue.

To evaluate the impact of surrounding genes on provirus expression, we turned to WGCNA, which reveals connections between genes that have correlated expression (Zhang & Horvath, 2005). This tool was used to create gene networks from the RNA-seq dataset of each body site. All expressed proviruses were analyzed alongside all expressed genes within 10kb of each provirus. We hypothesized that proviruses driven by read-through transcription would be strongly correlated with the surrounding gene that was driving this expression. Only one gene-provirus connection was significant, with the provirus 11q12.3 showing a significant connection with the gene ASRGL1 (Supplemental Figure 3). This provirus sits inside an intron of ASRGL1, which makes it a likely candidate for read-through transcription. Yet, even though 15 of the 38 expressed proviruses are similarly genic, this was the only provirus whose expression was significantly correlated with that of a surrounding or neighboring gene.

### HML-2 provirus transcription is affected by LTR sequence

The HML-2 proviruses in the human genome comprise three subtypes, named for LTR differences, but with a variety of defining mutations across the whole sequence that have accumulated over evolutionary time. For each transcribed provirus, it is important to understand both the drivers and the downstream effects of expression. The first characteristic we used to identify and group the proviruses expressed in the GTEx dataset was LTR subtype, which is a useful proxy for understanding sequence diversity and provirus integration age (Subramanian et al., 2011). The expression patterns associated with the different subtypes (LTR5A, LTR5B, and LTR5HS) are shown by the color coding (red, green, and blue, respectively) in Figure 3A. A neighbor-joining tree was generated using the 5’ and 3’ LTRs, when present, from all expressed HML-2 proviruses in the GTEx dataset (Fig 5A). All three LTR subtypes are represented in this group, which includes 18 LTR5HS proviruses, 7 LTR5A proviruses, and 13 LTR5B proviruses, comprising 40%, 33% and 56% of each 2-LTR HML-2 subtype, respectively. As expected (Johnson & Coffin, 1999), in most cases, LTRs flanking the same provirus are nearest neighbors on the tree, with the branch lengths separating them reflecting the time since integration. The few exceptions to this pattern, such as the proviruses at 3p25.3 and 19q13.12 reflect ancestral rearrangement mediated by recombination between proviruses at different chromosomal locations (Hughes & Coffin, 2001).

**Figure 5.**
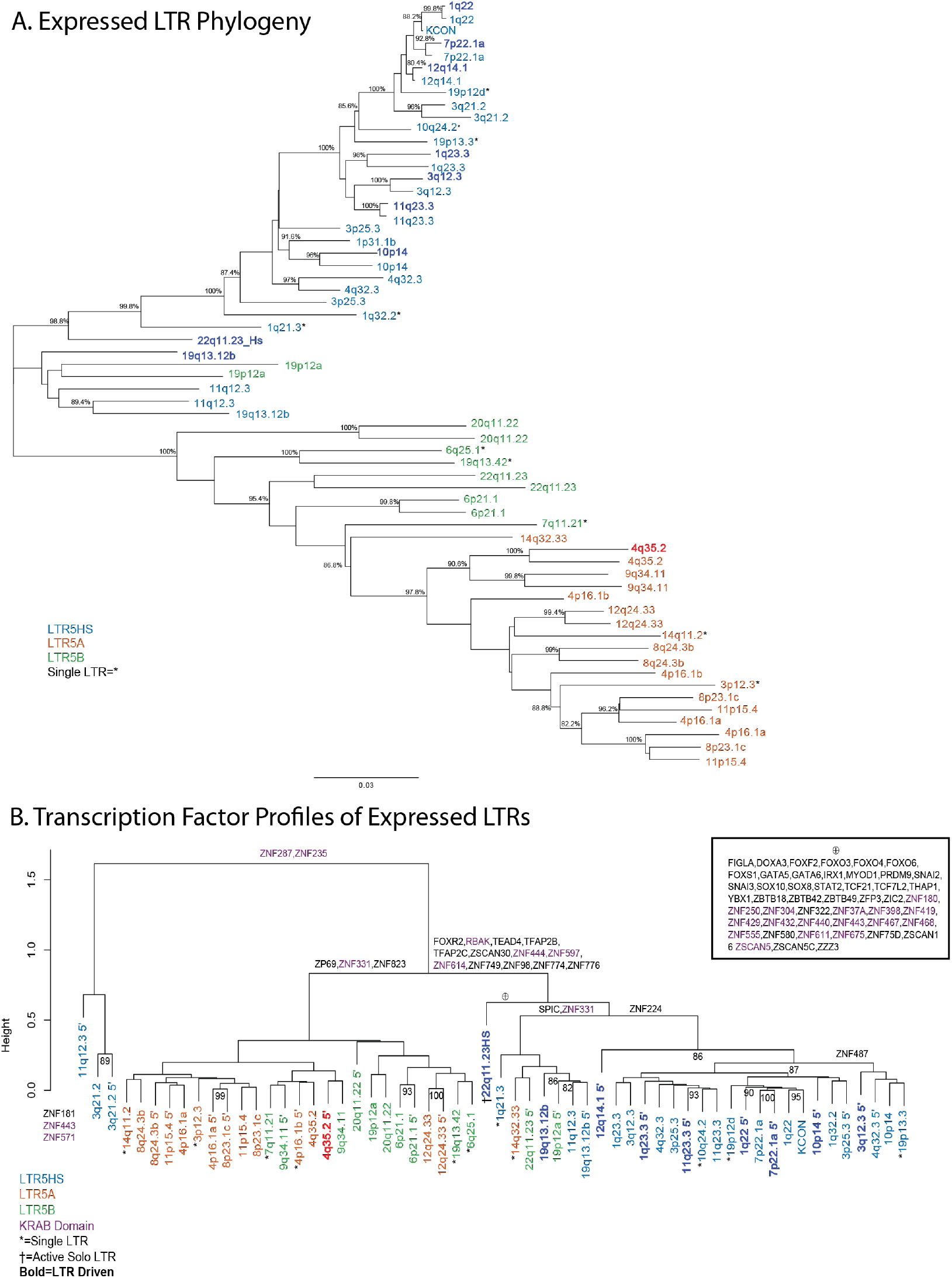
Relationships among multiple HML-2 proviruses of different LTR subtype. A. Neighbor-joining tree of the LTRs of each expressed HML-2 provirus in the GTEx dataset. Both LTRs are included when present, if only one LTR is present it is labeled with *. Color defines the subtype of each LTR as in Figures 1B and 3B. Darker color indicates proviruses whose expression is driven by the 5’ LTR. B. Cluster dendrogram of expressed HML-2 provirus LTRs based on TF binding profile (See Materials and Methods). Solo LTRs are denoted by *. LTRs are colored by LTR5 subtype. A darker color and **boldface** signify an LTR observed to drive provirus expression. Defining TFs are shown above the branches of the dendrogram. Purple TFs are those that are known to have a KRAB domain (Huntley et al., 2006). The large number of TF motifs that define the branch containing the 22q11.23 LTR5HS solo LTR, which drives the expression of the adjacent LTR5B provirus are shown in the inset under the ⊕ symbol.

As indicated by the greater p distance between their LTRs, LTR5A and 5B proviruses are the older groups, thought to have been active about ∼15-21 million years ago and to have gone extinct prior to the hominid divergence (Subramanian et al., 2011). None of the proviruses in these 2 groups contain an intact ORF for any viral gene (6 contain partial *gag* ORFs and one contains a polymorphic *gag* ORF, see Fig 8B).

One way to analyze mechanisms of provirus expression is to investigate the transcription factor binding motifs present in each LTR. The U3 region of the LTR includes a number of elements required for provirus expression including promoter and enhancer regions, which contain binding sites for transcription factors and other regulatory proteins (Manghera & Douville, 2013). The ability of transcription factors to bind to their cognate motifs in these regions can be modified by sequence variation and therefore this variation can have significant effects on provirus expression *in vitro* (Montesion et al., 2018b). To test whether the LTR sequence variation affected potential transcription factor binding and was therefore likely influence the pattern of HML-2 expression seen in the GTEx dataset, an alignment of expressed provirus LTR sequences was analyzed using the bioinformatics software FIMO to scan the DNA sequence of each provirus LTR for confirmed binding motifs of known human transcription factors (Grant et al., 2011). The LTRs were then clustered based on transcription factor motif presence this information (Figure 5B). This analysis separated the proviruses into two major clades, based on the presence/absence of ∼16 different transcription factor motifs with LTR5HS proviruses largely separating into one clade and most of the LTR5A and LTR5B proviruses combined in the other; the 5’ LTR of 11q12.3 along with both LTRs of 3q21.2 separated from the rest of the proviruses due to the absence of binding sites for ZNF287 and ZNF235, two factors with a currently unknown function. Notably, the 3’ LTR of 11q12.3 did contain the two ZNF TF binding sites and clustered with other LTR5HS and LTR5B proviruses. The LTR5HS containing clade was defined by the presence of 13 different TF motifs not found in the LTR5A/B clade. A number of these TFs were either involved in development (TEAD4, TFAP2B,TFAP2C) (Pastor et al., 2018; Satoda et al., 2000; Zhao et al., 2008) or known to be repressive (RBAK,ZNF597,ZNF614) (Huntley et al., 2006; Skapek et al., 2000). Interestingly, the 22q11.23HS solo LTR that drives the expression of the 22q11.23 provirus contains binding sites for 48 TFs that were not seen in any other LTR analyzed (Figure 5B, inset). This LTR is quite diverged from the other LTR5HS LTRs, sitting basal to nearly every other LTR5HS LTR besides 19q13.12b. The high-level clustering by transcription factor binding so closely replicating the clustering by LTR subtype suggests that the defining mutations of LTR5HS provirus LTRs have (or had) a significant effect on the ability of these proviruses to bind transcription factors and the consequent tissue specificity of expression. For example, the LTR 5A cluster on the left, from 14q11.2 to 8p21.3c has 5 provirus with 5’ LTRs with CNS (brain and cerebellum)-specific expression, and 12 of the proviruses in the 5Hs cluster from 1q23.3 to 11q 23.3 with 5’ LTRs show reproductive tissue specific expression. Interestingly, LTR5A/B LTRS appear to drive only one of the LTR-expressed proviruses while LTR5HS LTRS drive the other 8. This observation suggests that the 3 development-associated TF’s present in the LTR5HS branch could play a role in driving this reproductive tissue expression.

### HML-2 provirus transcription is biased towards older proviruses

HML-2 proviruses began accumulating in the primate ancestral genome around 35 million years ago and continued to accumulate in human ancestors in waves until less than 500,000 years ago (Figure 1A). This wide range of integration timing has created a diverse population of HML-2 proviruses in the human genome, each of which has been subject to unique selection pressure since integration. To understand the influence of proviral age on expression of each provirus in the GTEx dataset, the proviruses were grouped by the **E**arliest **S**hared **A**ncestor (**ESA**) of humans and modern primate species, creating 7 different age groups of proviruses (Rhesus Macaque, Gibbon, Orangutan, Gorilla, Chimpanzee, Human Specific-Fixed, Human Specific-Polymorphic). The expression of these groups was then broken down in two different ways: by the average expression of each group across all tissues tested (Figure 6A) and by the percentage of proviruses in each group that were expressed > 1 TPM (Figure 6B). Both methods revealed that, among the Hominoid-specific proviruses the “older” ones (ESA from Gibbon to Gorilla) were expressed to a higher level than the recently integrated ones shared with Chimpanzee or Human specific. Expression from proviruses that are shared with Gibbons makes up the majority of HML-2 transcripts in each tissue, along with proviruses that are shared with Gorilla (Figure 6B). The Orangutan group proviruses are the largest ESA group (11), expressed to one of the highest levels apart from Gibbon in Figure 6A, yet displaying a lower average when normalized to ESA group size in Figure 6B.The “younger” proviruses in the Chimpanzee and Human Specific groups were more highly expressed in certain tissues, including thyroid and testis. The level of expression of Human Specific-Polymorphic proviruses was also relatively higher in the thyroid and testis.

**Figure 6.**
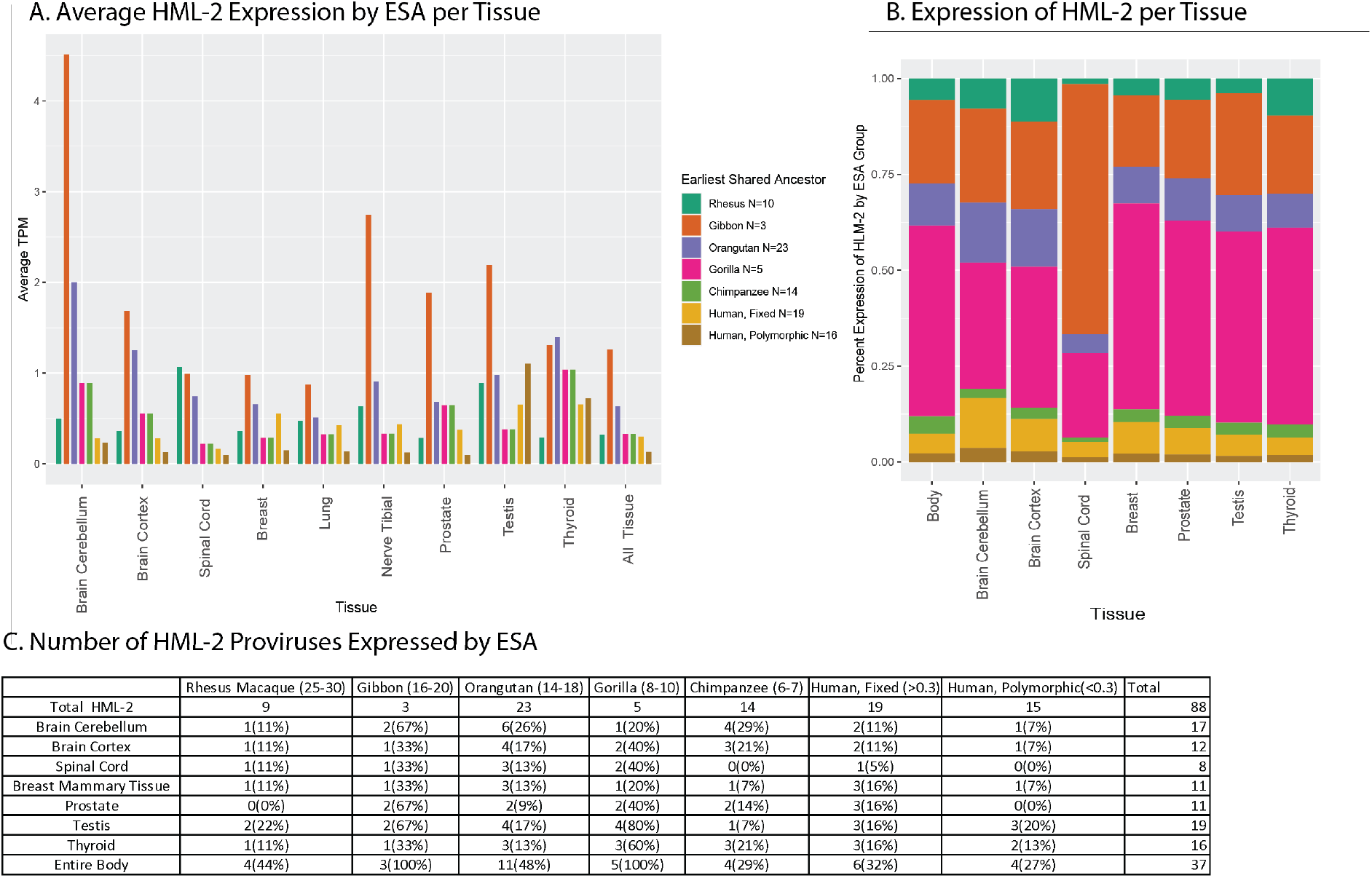
Expression of HML-2 proviruses in the GTEx dataset as a function of provirus age. A. The plot displays the average TPM of proviruses in the GTEx dataset grouped by the earliest shared ancestor (ESA). Proviruses are sorted by identification of orthologous insertions in species related to humans (Subramanian et al., 2011) and their average expression is displayed for the body sites of interest. B. The average expression of HML-2 proviruses in each ESA group normalized for the number of proviruses in that group and displayed as the percentage of total HML-2 expression at each body site. C. HML-2 expression >1 TPM broken down by ESA group. Numbers in parentheses next to the species names indicate estimated time, in millions of years, to their last common ancestor with humans.

Along with higher expression levels in specific tissues, the percentage of older proviruses widely expressed in most tissues was higher than that of the younger proviruses (Figure 6C). The cerebellum, testis and thyroid are the only tissues that contained a higher number of expressed Chimpanzee group and Human Specific group proviruses. These tissues appeared to promote expression from more of the younger proviruses as compared to other tissues. These data suggest that proviruses of different ESA groups are under different levels of control in the human genome where older proviruses that may have undergone longer periods of selection are more likely to be expressed. It is therefore possible that certain expressed proviruses might have provided a selective advantage that contributed to their fixation and protected them from being lost by drift or solo LTR formation.

### Covariates, including age and sex of donor, only slightly affect HERV-K(HML-2) transcription

The individual variation in expression of distinct proviruses across different tissues implies that not all non-diseased individuals express HML-2 proviruses in the same way. To address possible underlying causes of this variability, we utilized the metadata provided with the GTEx cohort to break down the donors according to different covariates. We hypothesize that inter-donor variation may be partially explained by the sex, age, and/or timing of death of the individual donors.

Biological sex can have an important impact on gene expression (Lopes-Ramos et al., 2020). We calculated total HML-2 expression in tissues of interest depending on sex as determined by the expression of Y chromosome genes (Figure 7A). At the total HML-2 level, no significant differences can be seen between the two groups in any tissue tested, with few exceptions such as breast and cerebellum, which displayed slight, but not statistically significant, differences in total HML-2 expression. Focusing specifically on these tissues, and the individual provirus expression in each of them, can offer more insight. In breast tissue samples, including 218 male and 134 female donors, 6 proviruses showed significantly higher expression in biological females than males (Figure 7D, p <0.05). Thus, presence or absence of Y chromosome gene expression may not correlate with overall HML-2 expression but can do so in certain body sites in a more specific and targeted way, although we cannot exclude sampling issues, such as distribution of specific cell types, that may affect the apparent differences in expression levels between the sexes.

**Figure 7.**
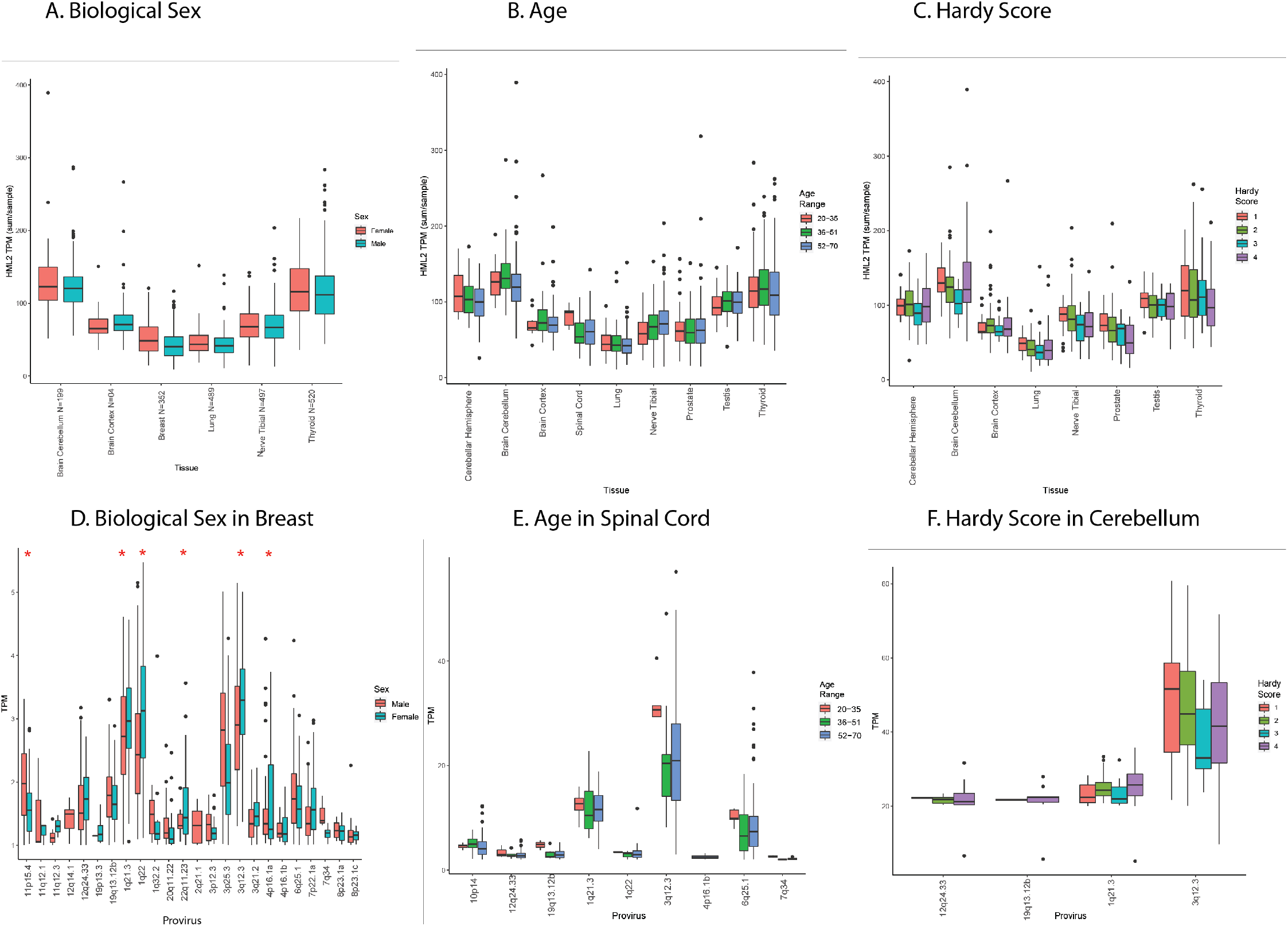
Effects of covariates and morbidities on HML-2 Expression. A,B,C. Average TPM per tissue of interest of total HML-2 expression for donors separated by Biological Sex, Age and Hardy Score plotted as in Figure 3A. Biological sex was identified by Y chromosome expression: age was divided into three groups (20-35, 35-51,and 52-70); Hardy score is broken down into 4 categories. 1 is the fastest death, 4 is the slowest. D,E,F. Expression of individual proviruses in the indicated body sites(Breast, Spinal Cord and Cerebellum). Average TPM of all proviruses expressed greater than or equal to a TPM of 1 is `displayed from all samples of each body site.. Red asterisks indicate statistically significant (p<0.5) differences (t-test).

The GTEx cohort includes donors ranging in age from 20 to 70 years, and therefore age-related characteristics could have affected HML-2 expression in donors to this dataset. To examine age-related changes in expression, samples for each tissue were sorted into three donor groups: ages 20-35, 36-51, and 52-70. Total HML-2 expression among the age groups was variable in certain body sites, particularly in the nervous system (Figure 7B). The cortex and spinal cord showed altered expression of HML-2 proviruses in the 20-35 age group (Cortex, N= 10 samples, spinal cord, N=55) as compared to older donors (cortex: 36-51 years, N=30; 52-70 years, N=164; spinal cord: 36-51 years, N= 23; 52-70 years, N=92), yet no differences were statistically significant. The lack of significance could be in part due to the small number of samples from the 20-35 age group (N=5). Alternatively, the cerebellum had lower expression of total HML-2 proviruses in the 52-70 age group (N=157) as compared to the younger groups (20-35 years, N=12, 52-70 years, N= 30). Much of this variation was the result of expression of the provirus at 3q12.3, which was elevated in the 20-35 age group in the spinal cord and had a large variation in expression in both the cortex and cerebellum (Figure 7E) resulting in the total HML-2 levels previously mentioned. The cerebellum also displayed a lower level of 1q21.3 expression in the 20-35 age group as compared to the 52-70 age group, a difference that was not observed in other body types. Similar to the effects of biological sex, the age of the donors appears to affect specific proviruses in individual body sites, and the provirus at 3q12.3 was the one whose expression was most affected by changes in age in multiple body sites measured.

Provirus expression could also have been artifactually affected by stress associated with the process of death. The Hardy score is a medical classification assigned by the GTEx consortium that describes how quickly an individual died, allowing one to take into account stress-related effects on gene transcription from post-mortem donors (Ferreira et al., 2018). Hardy scores are on a scale from 1 to 4, where 1 is assigned to death from accident or blunt force trauma that lasts less than 10 minutes, 2 is assigned sudden death of a previously healthy individual (e.g. myocardial infarction), 3 is assigned to a death over the course of 1 to 24 hours, and 4 is assigned to a death from a long-term illness. A quick violent death as compared to a longer one can have different effects on gene expression in sites across the body and was therefore an important effect to look at (Ferreira et al., 2018). On the level of total HML-2 expression the cerebellum showed unique patterns across the 4 Hardy scores (Figure 7C). In the cerebellum, HML-2 expression also decreased as the Hardy score increased until the 4th category where it increased again. This change was the result of increased expression in the 4th category of donors by the proviruses 3q12.3 and 1q21.3, which decreased over the initial 3 categories in the cerebellum (Figure 7F). The change in HML-2 expression as a result of cause of death was more uniform than the other covariates analyzed, with HML-2 expression the highest in the violent death category and decreasing as the length of the terminal phase increased. Yet it appears to rebound in the cerebellum in the 4th category. This observation could suggest retained activity in the brain for the long-term deaths that is not retained in other body sites. Overall, the covariates of biological sex, age and Hardy score can partially explain the variation in expression of certain proviruses in specific tissues. However, even among donors with similar covariates, the provirus expression can vary dramatically suggesting that more complex correlates of expression remain to be found.

### Potential for HML-2 protein expression

The HERV-K(HML-2) group is unique among HERVs in the number of proviruses that retain intact ORFs for viral genes. Products of these viral genes have been associated with a number of diseases and also could be providing beneficial effects, for example through viral restriction or immune modulation. While specific increases in some of these viral proteins have been observed in certain disease states, leading to suggestions that they might be useful as diagnostic indicators or therapeutic targets (Curty et al., 2020), the extent to which they are expressed across a non-diseased human body is largely unknown. In order to answer this question, we examined the sequence of each expressed HML-2 provirus, analyzing each viral gene (i.e., *gag, pro, pol, env, rec*, and the product of the type 1 mutant transcript *np9*) for the presence of deletions, stop codons, or frame shifts that would disrupt the open reading frame. Although it was not possible to evaluate the functionality of each gene, its ability to express a viral antigen could be inferred (Figure 8A). Across all HML-2 proviruses, intact ORFs are maintained for all viral genes. 11 proviruses contain an intact *gag* gene; 19 others can only encode one subunit of *gag*. 11 proviruses contain an intact *pro*, yet only 6 of these proviruses have an intact *gag* upstream for proper translation. 9 carry an intact *pol* gene, although only 3 proviruses possess an intact *gag* and *pro* upstream of the pol gene. Additionally, there are 8 proviruses potentially capable of expressing an intact *env* gene.

**Figure 8.**
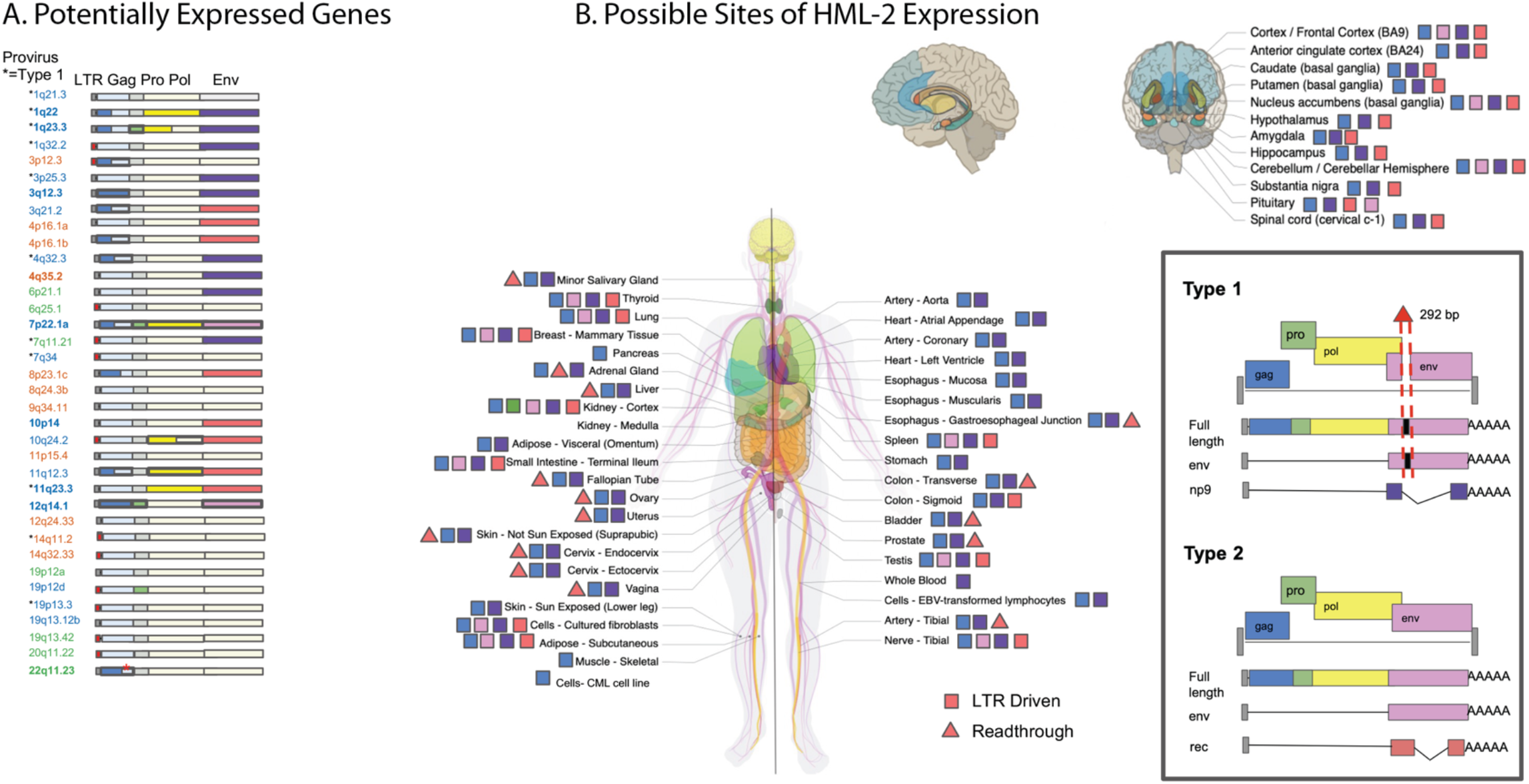
Potential consequences of HML-2 Expression in the GTEx dataset. A. Schematic of each provirus expressed at TPM > 1 in the GTEx dataset. Each ORF and LTR is designated with a colored box; a filled box represents an intact ORF. A partial LTR is represented by a red box. Colors are explained in the legend. Asterisks next to the provirus locations indicate type 1 proviruses, which are incapable of expressing *env* or *rec*, but rather express *np9*. Bold text represents a LTR driven provirus. B: Each body site included in the GTEx dataset is represented in this diagram. All non-brain tissues are labeled on the body; a zoom out of the brain labels each of its tissues individually. A colored symbol next to each body site denotes which intact ORFs could be expressed in that body site based on provirus sequence and transcriptome analysis. A square indicates LTR-expressed genes; a triangle indicates other potentially expressed genes. The box in the lower right displays the colors of each ORF and shows the differences between type 1 and type 2 proviruses. The * marks a polymorphic insertion in the provirus at 22q11.23 that breaks the Gag ORF in 43% of the population and is found in the hg38 reference genome.

Our large dataset of HML-2 expression data provided an opportunity to predict the pattern of potentially expressed viral proteins in different tissues. While the RNA-seq read length limits the identification of gene transcripts themselves, we could associate expression of each specific provirus with its intact gene content inferred from its DNA sequence (Figure 8A). These data were used to create the map seen in Figure 8B. This analysis reveals that very few of the expressed proviruses contain intact ORFs. There are 2 intact *gag* genes, coming from the proviruses 3q12.3 and 12q14.1. The provirus 3q12.3 is expressed in nearly every GTEx tissue, while 12q14.1 is expressed only in the kidney. None of the 3 known proviruses with intact *gag-pro-pol* sequences are expressed in GTEx tissue samples, although the kidney-expressed provirus at 12q14.1 contains an intact *gag*-*pro* sequence. Additionally, 4 different expressed proviruses contain individual intact *pro* or *pol* genes without a fully intact Gag polyprotein. Two proviruses, 7p22.1a and 12q14.1, express an intact env ORF, yet 7p22.1a is the only one with a functional *env* gene (Dewannieux et al., 2005). This provirus is expressed in 12 different tissues from numerous different organ systems. Despite there being only 2 intact Env ORFs expressed in the GTEx cohort, 6 different proviruses with an intact Rec ORF are expressed. These proviruses are expressed in body sites from multiple organ systems, suggesting that HML-2 Rec expression may be a common occurrence in non-diseased tissue. These data suggest that even in non-diseased individuals, HML-2 Gag can be present in as many as 53/54 body sites sampled and HML-2 Env could be present in 13/54 body sites. Therefore, the presence of HML-2 transcripts and possibly proteins, can be viewed as a normal part of the proteome and transcriptome of non-diseased tissue.

## Discussion

The role of HERVs in human biology is still largely a mystery. While the activity of recently integrated HML-2 proviruses has been previously studied in a number of disease contexts, their activity in non-diseased tissue has been largely unexplored. In this study we leveraged the scale of the NIH GTEx project to analyze HML-2 transcription in 54 different body sites from over 948 different donors. We detected HML-2 expression across the non-diseased human body, finding that 37 different proviruses are expressed and that every tissue site shows some level of expression. The pattern of expression is heterogenous: with many proviruses, it is likely affected by LTR sequence, at least partly through the unique set of transcription factor binding motifs contained in each LTR; with others, particularly those lacking 5’ LTR sequences, it is related to expression of nearby or surrounding genes. The considerable heterogeneity of expression of individual proviruses among the donors sampled, for the most part, remains unexplained, although biological sex also appears to significantly affect the expression of some proviruses in specific tissues. The overall HML-2 expression profile is largely made up of transcripts of evolutionarily older proviruses, yet some younger proviruses with intact ORFs are also expressed.

The highest total levels of HML-2 expression were found in the cerebellum, testis, thyroid and pituitary gland. The cerebellum and testis also supported the widest range of provirus expression of any tissue, with 17 and 19 proviruses expressed respectively. The varied pattern of HML-2 expression across tissues suggests the existence of tissue-specific factors that could drive this expression or certain tissues that exhibit a reduced restriction of provirus expression. In at least some cases, the HML-2 expression observed could be the result of cellular environments that promote widespread gene expression including many HML-2 proviruses. It is known that both cerebellum and testis express large numbers of tissue-enriched genes (van de Lagemaat et al., 2006). In other hotspot tissues like the thyroid and pituitary gland, the proviral expression observed could be a result of HML-2 response to different signaling hormones. HERVK LTRs are known to contain binding motifs for signaling hormones such as androgen, estrogen and progesterone which could have an activating role in these tissues (Manghera & Douville, 2013).Of course, the function of these predicted sites also depends on the expression of TFs in the relevant cell type, as well as other epigenetic features, including DNA, histone, and TF modifications, presence of negative TF elements, and more. More study such as ChIP-seq and functional analyses, will be required to identify the relevant factors and to confirm the activities predicted by sequence alone. Published data report that the factors SP1,SP3 and YY1 are involved in HML-2 LTR expression (Fuchs et al., 2011; Knossl et al., 1999) yet no motif for these factors was identified in this analysis. Additionally, it is unclear from our data what role repressive factors play in regulating HERV expression. Zinc finger proteins are known regulators of HERVs, via their ability to identify and bind motifs in the LTR and the PBS (Iouranova et al., 2022; Wolf et al., 2015; Yang et al., 2022). The KRAB-ZFPs have been reported to regulate DNA methylation and histone modification of HERVs through the recruitment of proteins such as KAP1/TRIM28 (Rowe et al., 2010). 86 such proteins were detected in our FIMO sequence analysis with 26 of them possessing KRAB domains (Huntley et al., 2006). It is likely that the activity of these proteins helps to regulate HERV expression in non-diseased tissue and the differential expression of these multitude of factors could result in the pattern of expression we observed.

It has been previously proposed that the expression of HERV sequences in the testis is a result of HERV involvement in development and reproduction(Larsson et al., 1994). According to our analysis, the LTR5HS subtype binds factors associated with the regulation of early development such as FOXR2 and TFAP2B suggesting that the expression in reproductive tissues like the testis could indeed have a regulatory role. We consider it more likely, however, that the HML-2 activity in reproductive tissue and the testis in particular could be a relic of ancient viral infection (Diehl et al., 2016). Although alternative mechanisms have been proposed, most evidence supports the interpretation that ERVs in general, and HML-2 in particular arose from infection of germ line cells with the corresponding exogenous virus, likely during epi- (or pan-) zootic infection in ancestral species (Bannert & Kurth, 2006). According to this view, at the time of integration, the LTR of each newly-minted provirus must have directed an expression pattern adapted to support a virus lifestyle, including replication in sites that promote transmission from one individual to another, be it sexual, blood contact, or vertical (mother to infant) (Diehl et al., 2016). With evolutionary time, these patterns can be altered by forces of mutation, selection, and drift acting on the proviruses or the individuals containing them, leading to loss, or inactivation of the proviruses whose expression is deleterious or even neutral and over long periods of evolution to exaptation to a new function. An example of the latter process may be the provirus at 3q12.3, an old LTR5HS type 1 provirus, whose ESA is gorilla, and whose LTR-driven expression is found in nearly all body sites examined (Figure 3A,, 4A). The 5’LTR of this provirus has a 4 bp duplication, which is fixed in the human population, but not present in gorillas or chimpanzees, and which creates a binding site for HOX-PBX family of transcription factors, likely responsible for the widespread expression (Montesion et al., 2018b). The predominant expression of younger LTR5HS proviruses in reproductive tissue may reflect persistence of the original viral expression pattern, while expression of the older LTR5A/B proviruses in CNS tissue, with no obvious relationship to viral replication or transmission, may reflect their post integration evolution to completely different functions, which remain to be discovered. Thus, the greater frequency of expression of older than younger proviruses might be due to selective protection of these proviruses from loss due to drift through (for example) solo LTR formation.

While many HML-2 proviruses have been damaged by mutation and deletion, 9 of the expressed proviruses contain an intact 5’ LTR exhibit and LTR-driven expression, placing an increased importance on the binding motifs in their 5’ LTRs. It is therefore interesting that all but one is an LTR5HS LTR. The observation that the regulatory and developmental motifs in the LTR5HS sequence directly regulate transcription of these proviruses and the role of LTR5HS expression in relation to development require further study.

The presence of intact LTR5HS LTRs that are actively driving transcription would seem to imply that the comparatively younger HML-2 proviruses are expressed to a higher degree than the older proviruses. Yet, HML-2 expression across the body is largely made up of older proviruses, mostly expressed via non-LTR mechanisms, to a higher level than younger proviruses. This selective expression is made more interesting when considering which proviruses contain intact viral ORFs. Only 2 of the 11 proviruses with an intact Gag are expressed in non-diseased tissues, as are 2 of the 8 proviruses with an intact Env, along with none of the proviruses with a fully intact Gag-Pro-Pol polypeptide found in the GTEx data. The proviruses with retained intact ORFs are generally younger than the more defective ones, reflecting the lesser accumulation of damaging mutations over time. Therefore, it appears that the bias of expression of older provirus genes is due to the lower concentration of intact ORFs in these insertions. They could be expressed due to adaptive mutations that drive some beneficial activity such as enhancing host gene expression, as proposed by Xiang et al (Xiang et al., 2022). Alternatively, the expression of these proviruses could be retained merely due to the lack of viral protein expression driving any purifying selection against them.

If the majority of HML-2 proviruses that contain intact ORFs are repressed to prevent the possible deleterious expression of viral proteins, the viral ORFs that are still expressed may well have a higher chance of having a beneficial effect on the host. As previously discussed, there are numerous examples of ERV *gag* and *env* genes being co-opted by their hosts for beneficial purposes, with defense against exogenous viruses being the most common function (Best et al., 1996; Blanco-Melo et al., 2017; Murcia et al., 2007; Robinson et al., 1981). Viral proteins could also be co-opted for another mechanistic benefit such as transport of signaling molecules in a Gag-like protein shell (Pastuzyn et al., 2018). All expressed proviruses observed in GTEx that contain intact ORFs have had to endure hundreds of thousands to millions of years of selection in the human ancestral genome and still retain the ability to be expressed and contain specific intact ORFs. 3 of these proviruses (3q12.3,7p22.1a and 12q14.1) are also LTR driven which leaves a viral ORF under direct LTR control. The provirus at 3q12.3 is expressed throughout the body at high levels and contains an intact *gag* gene, making it a prime candidate for co-option that is worthy of future study. 12q14.1 is a provirus that retains both an intact *gag* and *env* ORF and yet it is only expressed in the kidney, possibly pointing to a much more targeted function inside that organ. 7p22.1a carries a well-known fully functional *env* gene that has been studied previously in the literature (Dewannieux et al., 2005). Together these ORF-containing proviruses are highly expressed and LTR driven, likely creating translatable viral transcripts in tissue throughout the body (as shown by squares in Figure 7B). By contrast, translation of the predicted ORFs of the non-LTR expressed proviruses depends on the structure of the individual transcripts (as shown by triangles in Figure 7B). Further analysis of the protein products of each of these proviruses could reveal a role for an HML-2 provirus in normal human biology, as has been previously uncovered for other HERV families (Lavialle et al., 2013).

The presence of proviral transcripts and the likely expression of viral proteins in tissues through the non-diseased human body reveal a constant expression of HML-2 proviruses, with potential implications for the clinic. Upregulated expression of HML-2 characteristic of some cancers, is being studied as both a phenotypic marker and an immunotherapeutic target for cancer detection and treatment (Curty et al., 2020). For example, The provirus at 12q14.1 is expressed uniquely in the kidney cortex (Figure 3B), and encodes an intact, albeit nonfunctional Env protein (Dewannieux et al., 2005). Therefore, clinical studies of antibodies or cytotoxic T-cells directed against HML-2 Env epitopes will need to take potential off target effects due to expression in normal tissue into account into consideration in their design. It will also be important to keep in mind that large differences in the nature, distribution, and expression patterns of ERVs from one species to another will make it impossible to test such safety issues in standard animal models.

While more work will be required to understand the mechanisms driving their expression and the differences in expression between diseased and non-diseased tissue, it will be important to remember that HML-2 transcripts can always be found in the human body. Therefore, using the level of provirus expression or the presence of viral protein as a phenotypic marker will require controlling for this activity of non-diseased expression of 37 HML-2 proviruses across the body. Our characterization of HML-2 expression in the GTEx database should therefore serve as a useful resource for the clinical application of HML-2 moving forward.

## Materials and Methods

### RNA-seq analysis of the GTEx cohort

The entire RNA-seq portion of the GTEx cohort V7 was downloaded from dbGaP under phs000424.v7.p2 (Mailman et al., 2007; Tryka et al., 2014) https://www.ncbi.nlm.nih.gov/projects/gap/cgi-bin/study.cgi?study_id=phs000424.v7.p2) in fall 2019. Cohort V8 was downloaded from AnVIL under phs000424.v8.p2 (https://anvil.terra.bio/#workspaces/anvil-datastor-age/AnVIL_GTEx_V8_hg38) between Fall 2020 and Spring 2021. The analytical workflow is outlined in Figure 2A. RNA-seq data were downloaded in SRA format and subsequently paired-end fastq format were extracted using the SRA Toolkit (Leinonen et al., 2011). V8 files were downloaded as aligned bam files and converted to fastq using a python script that calls Picard samtofastq provided by GTEx (Consortium, 2020). Resulting fastqs were passed to FastQC to check initial read quality, and files were passed to Trimmomatic to remove reads that fell below our threshold (phred score < 30, length < 75 bp, adapter sequence presence). Reads were aligned to hg38 (UCSC version, 4) using HISAT2 (Kim et al., 2019). Alignment files were passed to Telescope (Bendall et al., 2019) to generate provirus-corrected counts files. Individual Telescope files are grouped by tissue using the “Telescope_Merge_Counts” script. V7 and V8 files had to be merged separately, due to the differences between the usage of SRA ID by dbGAP and SUBJID by AnVIL. Separate R scripts were utilized to address this matter. Raw counts were TPM normalized in R prior to any further analysis. All code is available at (https://github.com/Coffinlab/GTEx_HML2).

### Expression Plotting

The boxplot of total HML-2 expression was generated using the script “HML2 expression-HML2 per tissue”. This script converts raw counts to TPM and then groups provirus counts per tissue. These counts are then averaged per sample number and plotted in the boxplot. The script “graph heatmap of individual HML2 proviruses” was used to generate the provirus heatmap. This script took the previously generated TPM counts and calculated the average TPM in each tissue for each provirus. These averages were then fed into pheatmap to generate the figure.

### Covariate Statistical Analysis

The statistical analysis completed on HML-2 counts for biological sex, age and Hardy score were completed in R using Limma and Voom. This analysis was completed using the script “individual provirus covariate plotting”. Each tissue’s raw counts were filtered for low-expressed genes and a multidimensional scaling plot was generated to check for proper clustering of samples. The raw counts were then fitted to a curve using Voom before limma was used to fit a linear model to the data and calculate contrasts and significance between each covariate group.

### LTR Phylogeny

2-LTR proviral HML-2 sequences were downloaded from the hg38 human sequence database on the UCSC Genome browser and loaded into BioEdit. This file was aligned to the KCON consensus sequence (Lee & Bieniasz, 2007) using the online MUSCLE interface through EMBL-EBI(Edgar, 2004; Madeira et al., 2019). The 5’ LTR: (corresponding to bp 1-968 of the consensus) of each provirus, and 3’ LTR: (bp 8505-9472), or 3’ LTR alone, when a 5’ LTR did not exist, were extracted from the total provirus genomes and realigned using MUSCLE. This alignment of LTR sequences was then loaded into MEGA X and a neighbor joining tree was constructed with 500 Bootstrap replications. The labels and colors of the tree were then edited in Figtree.

### Transcription Factor Binding Motif Identification

The HML-2 LTR alignment, curated for the phylogeny, was run through the FIMO software (Grant et al., 2011). This software searches each sequence for individual matches of provided motifs. We provided FIMO the binding motifs of all human transcription factors from CIS-BP (Weirauch et al., 2014) along with an alignment of each 5’ and 3’ HML-2 LTR. The p value threshold used was 10^−4^. The R script “TF Motif dendro solo” was then used. This script loads the .tsv file of identified motifs, filters out unused LTRs and creates a matrix of motifs per LTR. This matrix is clustered using pvclust (ward.D2 method and 1000 bootstrap value).This clustering is then plotted before labels and colors are added in Adobe Acrobat.

### Provirus Age Analysis

HML-2 proviruses were grouped by the earliest shared ancestor(ESA) for each provirus by comparing orthologous insertions in related species (Subramanian et al., 2011). These groups were then used to calculate average expression in each tissue for Figure 6A. Each group average was then scaled by the number of proviruses in each group and plotted as a percentage of HML-2 expression in Figure 6B. This data were then used to generate the two bar plots shown. The analysis was completed using the script “Provirus Age Expression”. The totals of expressed proviruses in each ESA group are displayed in Figure 6C.

### ORF Determination

HML-2 proviruses were aligned to the full length HERV-K (HML-2) provirus, 19p12b, with MUSCLE. Proviral segments aligned to 19p12b *gag* (nt 1112-3112), *pro* (nt 2938-3918), *pol* (nt 3915-6749), *env* (nt 6451-8550), *rec* (nt 6451-6711/8411-8467), and *np9* (nt 6451-6494/8411-8591) were translated with ExPASY (https://web.expasy.org/translate). Viral ORFs which lacked nonsense mutations and frameshifting indels were run through Motif Scan (https://myhits.sib.swiss/cgi-bin/motif_scan). Only ORFs that retained all predicted Pfam domains were determined to be intact. Additionally, *pro* and *pol* genes were only considered intact if they remained translatable in the context of the GagProPol polypeptide, as a deleterious N-terminal mutation should ablate translation of downstream ORFs.

## Acknowledgments

We thank Rebecca Batorsky for assistance with the bioinformatics analysis and William Johnson for help with the FIMO analysis.

This work was supported by Research Grant R35 CA200421 from the National Cancer Institute.

## Supplemental Table 1

**Table S1.**
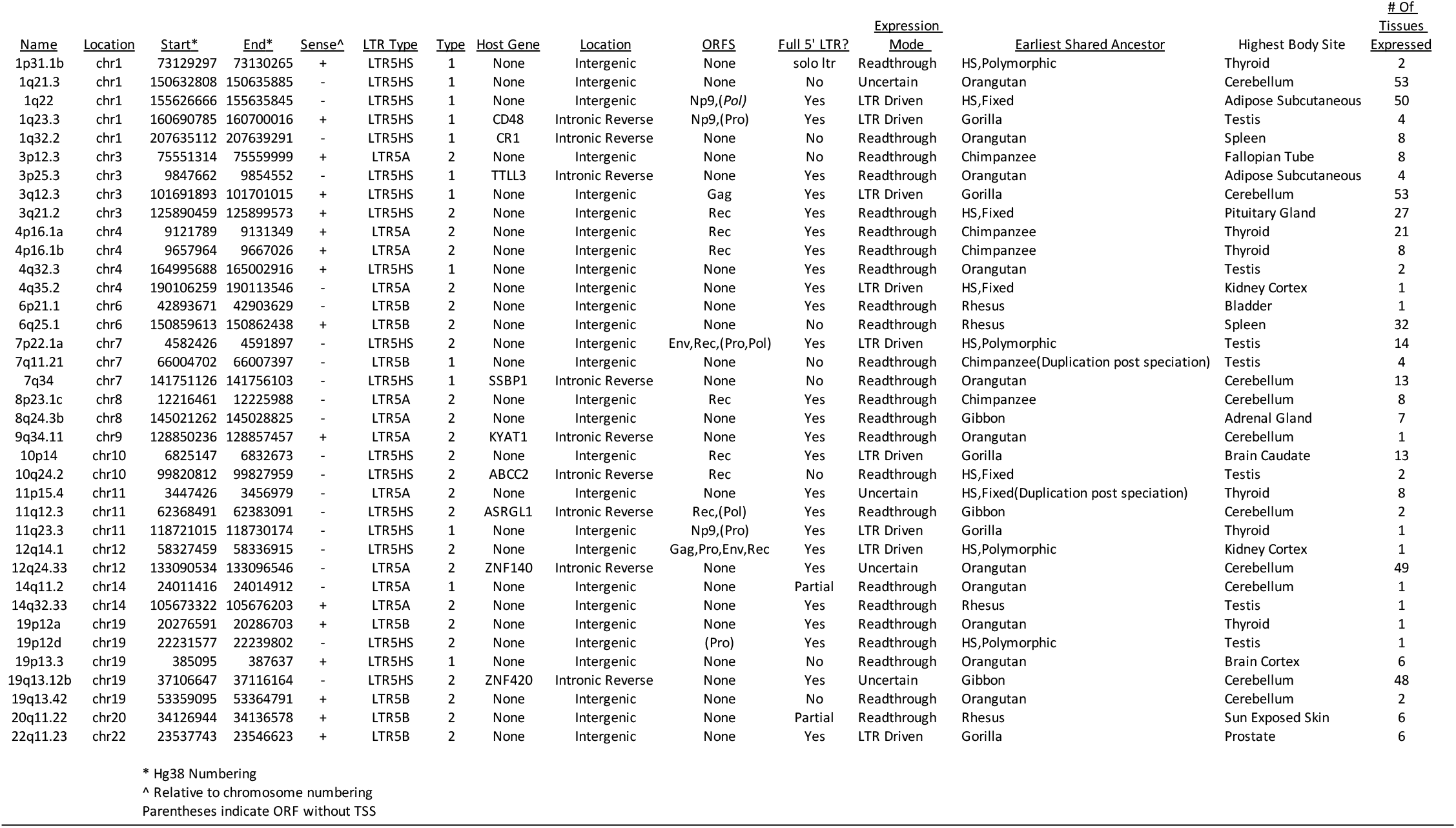
Properties of HML-2 Proviruses Expressed in the GTEX Dataset

## Supplemental Figures

**Supplemental Figure 1.**
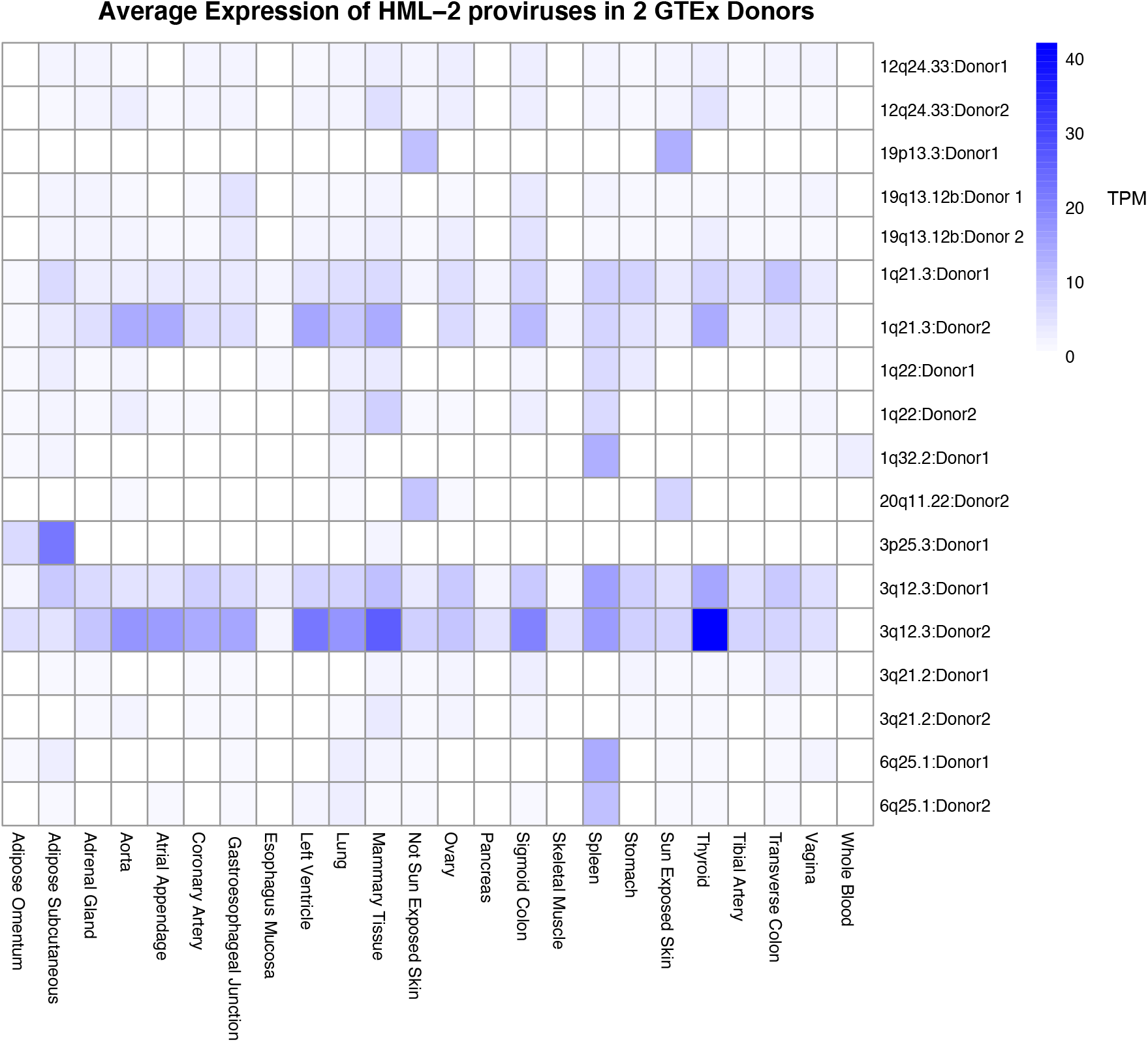
Comparison of individual donors This figure displays a heatmap of individual HML-2 expression from two female GTEx donors. Provirus expression is given in TPM for each provirus detected (10 in Donor 1 and 8 in Donor 2) for 22 body sites. Each provirus is labeled on the side with which donor it was measured in.

**Supplemental Figure 2.**
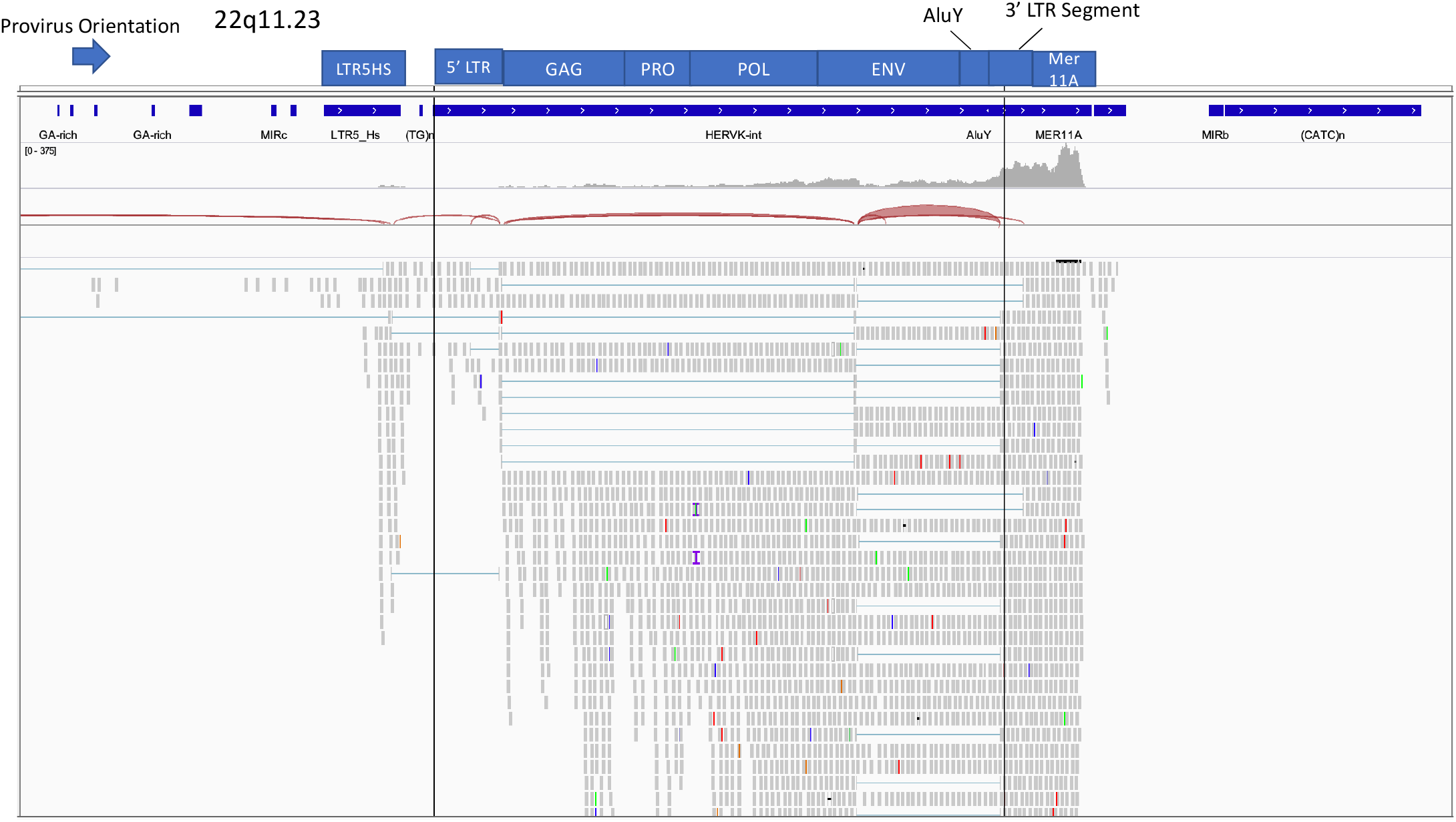
22q11.23 IGV Screenshot This figure displays the alignment of RNA-seq reads to the provirus 22q11.23 in a prostate sample visualized in Integrated Genomics Viewer. The sequence of 22q11.23 is defined by the vertical black lines. The LTR5HS LTR is displayed upstream to the left of the black line. The blue bars shown above the image indicate repeat elements as defined by the Repeatmasker track for HG38 downloaded from UCSC.

**Supplemental Figure 3.**
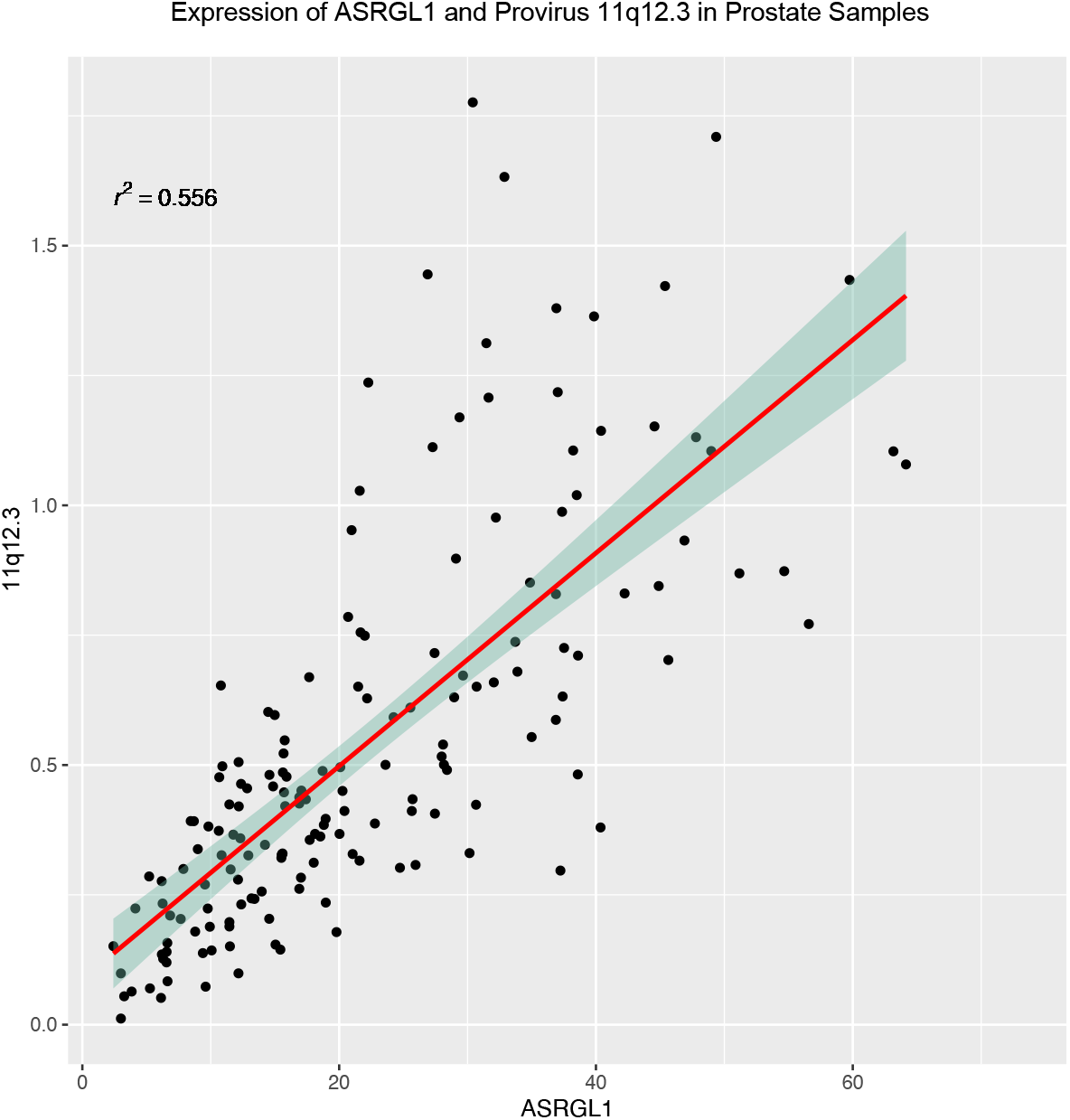
Scatterplot of ASRGL1 and 11q12.3 in Prostate tissue This figure displays a scatterplot of expression of both 11q12.3 and ASRGL1 expression in 166 Prostate samples in the GTEx dataset. Outliers more than 1.5 times beyond the upper and lower quartiles were removed. Smoothing line was added using linear model method in R ggplot2. A 0.95 confidence interval is displayed around the line.

